# Identification of Significant Gene Expression Changes in Multiple Perturbation Experiments using Knockoffs

**DOI:** 10.1101/2021.10.18.464822

**Authors:** Tingting Zhao, Guangyu Zhu, Patrick Flaherty

## Abstract

**Motivation:** Large-scale multiple perturbation experiments have the potential to reveal a more detailed understanding of the molecular pathways that respond to genetic and environmental changes. A key question in these studies is which gene expression changes are important for the response to the perturbation.

**Results:** We present here a method based on the model-X knockoffs framework to identify significant gene expression changes in multiple perturbation experiments. This approach makes no assumptions on the functional form of the dependence between the responses and the perturbations and provides finite sample false discovery rate control for the set of important gene expression responses. In a large-scale multiple perturbation gene expression data set from the Library of Integrated Network-Based Cellular Signature (LINCS) NIH program, we identified important genes whose expression is modulated in response to perturbation with anthracycline, vorinostat, trichostatin-a, geldanamycin, and sirolimus. Furthermore, we compared the set of important genes that respond to these small molecules to identify co-responsive pathways.

**Availability and Implementation:** https://github.com/flahertylab/deepYknockoff

**Supplementary information:** Supplementary data are available at *Bioinformatics* online.

## 1 Introduction

The elucidation of the mechanisms underlying cellular function requires perturbation of the system [Subramanian et al., 2017]. Perturbation approaches have been successful in understanding fundamental pathways in yeast [Hillenmeyer et al., 2008], humans [Shim et al., 2017], and other organisms [Skerker et al., 2013]. Perturbation experiments also provide a deeper understanding of the mechanism of action of small molecule compounds and guide opportunities for drug repurposing [Stathias et al., 2018].

A key bioinformatic step in the analysis of multiple perturbation experiments is to identify which genes are required for the adaptation to the perturbations. In a statistical modeling framework, the perturbation is the explanatory variable and the transcriptional changes of each gene are the response variables. Our objective is to identify those genes whose transcriptional response is associated with perturbations even after accounting for all the other genes. Solving this problem enables us to better understand the mechanisms of adaptation to environmental changes which in turn improves our understanding of human disease and helps us develop novel therapies [Keenan et al., 2018]. To address these questions, we formulate the following response selection problem.

### Problem Statement

Let *X_i_* encode the *i*-th perturbation and let *Y_i_* encode the gene expression measurement vector in response to the *i*-th perturbation. Assume we have *n* i.i.d. random variables (*X_i_, Y_i_*), where *X_i_* ∈ ℝ^*p*^, *Y_i_* ∈ ℝ^*r*^ assembled into two data matrices **X** ∈ ℝ^*n*×*p*^ and **Y** ∈ ℝ^*n*×*r*^ such that the *i*-th row of **X** is *X_i_* and the *i*-th row of **Y** is *Y_i_*. For example, in multiple perturbation experiments, *X_i_* is an indicator vector for the perturbation over *p* possible perturbations, and *Y_i_* is the *r*-dimensional transcriptional profile. A response variable *Y_j_* for *j* ∈ {1,…, *r*} is said to be unimportant if and only if *Y_j_* is independent of *X* conditionally on the other responses *Y*_−*j*_, where *Y*_−*j*_ = {*Y*_1_,…, *Y_r_*}\{*Y_j_*}. The set of unimportant variable indices is denoted by 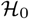 and we call a variable *Y_j_* important if 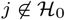. The set of the important variable indices is denoted as 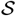. The unimportant responses are conditionally independent of the covariates given the important responses 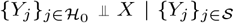. Our goal is to identify as many important responses as possible while keeping the false discovery rate (FDR) under control, where 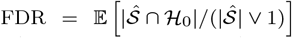 and 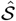 denotes a selected subset of the important response variable indices.

Despite the prevalence and importance of response selection problems, the literature on response selection methods is limited. The few methods that exist are restricted to the linear regression setting [An and Zhang, 2017, Su et al., 2016]. Moreover, these methods do not provide guarantees on the accuracy of the selected set with a finite sample size. Thus, the current field lacks both a general framework and concrete techniques to perform high quality response selection. To ensure selection quality, it motivates the need to control the expected fraction of false discoveries in response selection problems [Benjamini and Hochberg, 1995]. Furthermore, response selection methods should handle both linear models and complex nonlinear relationships between the responses and the features. To fill this gap, we take inspirations from a recent novel controlled feature selection method model-X knockoffs [Candes et al., 2018].

### 1.1 Review of Model-X Knockoffs

Model-X knockoffs generate knockoff features that act as negative controls in feature selection problems. These knockoff features mimic the dependence structure among the original features, but are independent of the response so that if a knockoff feature is selected by a feature selection procedure, it is known to be a false positive. The proportion of knockoffs selected by a feature selection procedure can be used as an estimate of FDR among the original features.

#### Definition 1

(Model-X Knockoffs [Candes et al., 2018]).Model-X knockoffs for the family of random variables *X* = (*X*_1_, …, *X*_*p*_) are a new family of random variables 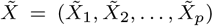 constructed with the following two properties:

1. For any subset 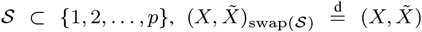: swapping the entries *X_j_* and 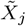 for each 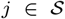 leaves the joint distribution invariant,
2. 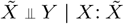 is independent of responses *Y* given the feature *X*.

Knockoff features constructed in this way have the following three properties for an explanatory variable *X_j_*, normalized to have zero mean and unit variance:

1. 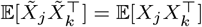 for *j* ≠ *k*,
2. 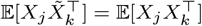 for *j* ≠ *k*, and
3. 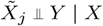, for all *j*.

These properties ensure that the knockoffs have the same correlation structure as the original variables, but are unrelated to the response by construction.

In general, constructing knockoff features that have these properties is challenging. One algorithm for generating knockoffs from an exactly known *F_X_*, sequentially from conditional distributions, has been suggested for a Gaussian distribution [Candes et al., 2018], a hidden Markov model [Sesia et al., 2019]. A Metropolis-Hastings formulation is applicable to all possible (but exactly known) *F_X_* [Bates et al., 2020]. To address situations where the marginal distribution is unknown, some works have proposed using deep generative models to learn a knockoff generating distribution [Jordon et al., 2018, Romano et al., 2020].

The general algorithm to perform controlled feature selection under the model-X knockoff framework has three steps. We first summarize the three key steps and provide more details about each step.

1. Step 1: Generate knockoff features 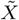 that satisfy Definition 1.
2. Step 2: Define the feature importance measure and the knockoff statistics for each *X_j_*, *j* ∈ [*p*], where [*p*] = {1, 2,…, *r*}.
3. Step 3: Decide the filtering threshold to guarantee the controlled FDR level.

#### Step 1: Knockoff Generation for Gaussian Features

Under the model-X knockoffs framework, it is assumed that the original features *X* come from a Gaussian distribution and the joint distribution for *X* and its knockoffs 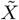 also follows a multivariate Gaussian distribution. If *F_X_* is a Gaussian distribution, properties of the multivariate Gaussian can be used to find the exact sampling distribution for knockoff generation. If 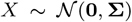, where **Σ** ∈ ℝ^*r*×*r*^, then a joint distribution obeying the pairwise exchangeability and conditional independence properties (Definition 1) is

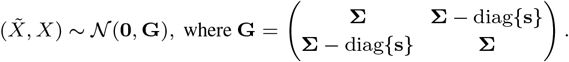

Since the conditional distribution of a multivariate Gaussian has a closed form, model-X knockoffs 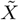 can be sampled from

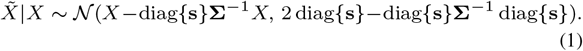

The value of diag{**s**} needs to be selected such that the joint covariance matrix **G** is positive definite and to ensure high power, larger values of **s** are preferred since 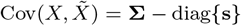. A detailed description of the procedure to select **s** is provided in Candes et al. [2018].

#### Step 2: Model-X Knockoff Statistic

Given **X**, **Y**, and 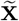 the generated knockoff features, the next step is to define and compute a feature knockoff statistic 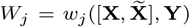 for each *X_j_, j* ∈ [*p*]. The response knockoff statistic *w_j_* must satisfy the following flip-sign property:

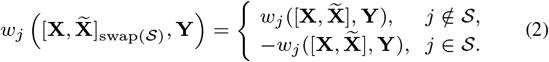

Constructing the knockoff statistics (*W*_1_,…, *W_r_*) has two steps. First, define the feature importance measures

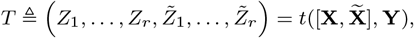

with 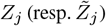 measuring the importance of 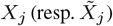. The feature importance measure must have the property that switching *X_j_* with 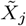 simply switches the component of *T* in the same way:

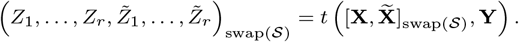

Second, the knockoff statistics constructed as 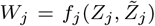 obeys the flip-sign condition (2), where *f_j_* is any anti-symmetric function.

#### Step 3: Filter with FDR control

The final step selects a set of *important features* 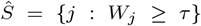. Model-X knockoffs have provided guidance on how to choose the threshold *τ* to ensure finite-sample controlled FDR. Let *q* ∈ [0, 1]. Given knockoffs statistics, *W*_1_,…, *W_r_* satisfying (2), let

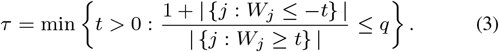

An appropriate procedure that selects the features 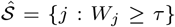 controls the FDR such that

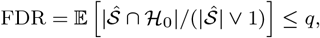

where 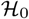 is the set of unimportant indices, 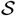 is the set of the important indices, and 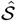 denotes a selected subset of the important indices. Since the procedures may seem abstract at first glance, we provide two concrete examples using Lasso and deep neural networks (DNNs) to illustrate the knockoff statistics construction in the context of response selection in Section 2.3 and 2.4.

### 1.2 Other Related Work

Many related methods have been proposed for identifying associations between an experimental perturbation and the transcriptional response of a single gene or a set of genes. But, these methods largely test different hypotheses than model-X knockoffs.

Traditional methods for variable selection, such as permutation testing or T-tests, focus on marginal testing. The hypothesis under consideration for these tests is *H*_0*j*_ : *X_j_* ⫫ *Y*. While marginal tests are a powerful exploratory data analysis tool, they may select variables that are not causally related to the response, but are instead only correlated with another explanatory variable. It is possible that conditioning other variables renders the features independent of the response.

Conditional tests seek to reject the null hypothesis 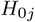 : 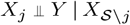, where 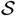 denotes the set of features that are *included in the model*. But the form of the relationship between the response *Y* and all the explanatory variables 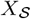 must be specified in the model; this includes all of the interactions between explanatory variables and the functional form of the relationship between those variables and the response.

In contrast, knockoffs seek to to test the hypothesis: *H*_0*j*_ : *X_j_* ⫫ *Y* | *X*_−*j*_. That is whether the explanatory variable is conditionally independent of *all of the other explanatory variables*, not just the set included in a generalized linear model. Furthermore, model-X knockoffs do not assume a functional form for the relationship between *Y* and *X* making them an appropriate tool for multiple perturbation experiments where the functional relationship between the dose of a perturbation and the transcriptional response is typically unknown and better left unspecified a priori. Finally, there has been a vast amount of research on feature selection in different settings [Fan and Lv, 2008, Tibshirani, 1996]. However, these methods, in general, do not control the FDR with finite-sample guarantees.

#### Summary and Contributions

This paper develops a methodology to identify important transcriptional changes in response to chemical and genetic perturbations by performing controlled response variable selection inspired by model-X knockoffs with theoretically guaranteed FDR control. We prove that the same properties that model-X knockoffs enjoy for feature variable selection hold for response variable selection problems due to a symmetry in the proof of the validity of the framework. We demonstrate a way to employ knockoffs for response variable selection in a linear model (Lasso) and we develop a way to employ knockoffs in a nonlinear model (DNNs) using a competing hidden layer. We analyze the L1000 phase I dataset from the NIH LINCS Consortium and identify important genes whose expression is modulated in response to perturbation with multiple small molecules including vorinostat, geldanamycin, sirolimus, trichostatin-a, wortmannin, and anthracycline.

## 2 Methods and Materials

In this section, we describe a method for response variable selection based on model-X knockoffs which we denote model-Y knockoffs. Section 2.1 provides a proof that model-Y knockoffs provide finite sample false discovery rate control for response variable selection. Section 2.2 presents an algorithm for generating and using model-Y knockoffs in a manner analogous to model-X knockoffs. Section 2.4 gives a non-trivial method for employing model-Y knockoffs for response variable selection in DNNs.

### 2.1 Model-Y Knockoffs

The key idea to adapting model-X knockoffs for response variable selection is the observation that the generation and use of model-X knockoffs only depends on the joint distribution between the explanatory variables and the response *F_XY_*. Model-X knockoffs assume that the marginal distribution *F_X_* is known exactly and this allows the conditional distribution *F*_*Y*|*X*_ to be unspecified. In response variable selection problems, we have abundant information about the marginal distribution *F_Y_*, so we use the fact that the joint distribution, can be factorized as *F_XY_* = *F*_*X*|*Y*_ *F_Y_*. This leads to a symmetry that carries through the theory that we exploit in our development of model-Y knockoffs.

#### Definition 2

(Model-Y Knockoffs). A random vector 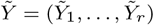 is the model-Y knockoffs of a random vector *Y* = (*Y*_1_,…, *Y_r_*) if it satisfies two properties: (1) Pairwise exchangeability: for any subset 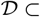 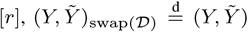, i.e. swapping the entries *Y_j_* and 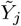 for each 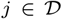 leaves the joint distribution invariant; and (2) Conditional independence: 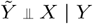, i.e., 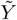 is independent of features *X* given the response *Y*, which can be guaranteed by constructing 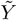 without looking at *X*.

Using this definition, we can prove the following theorem establishing the finite sample false discovery rate control for response variables.

#### Theorem 1.

Define the joint distribution of *X* and *Y* as *F_XY_*, the marginal distribution of *Y* as *F_Y_* and the conditional distribution as *F*_*X*|*Y*_. We assume that *F_Y_* is specified and *F*_*X*|*Y*_ is unconstrained. If 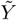 is the model-Y knockoffs of a random vector *Y* = (*Y*_1_, *Y*_2_,…, *Y_r_*) satisfying Definition 2 and *X* is the features such that 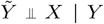, by swapping *X* and *Y*, the swapped 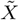 (which is original 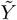 before swapping) is the model-X knockoffs of *X* (which is original *Y* before swapping).

Proof. We observe that the joint distribution of *X* and *Y* can be factorized as

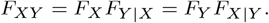

According to Definition 1, we have that for any subset 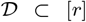, 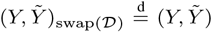, i.e. swapping the entries *Y_j_* and 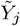 for each 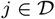. Once we swap *X* and *Y*, we have that 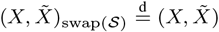, where 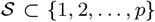 and we set *p* = *r* such that 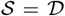. We also have that 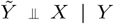, which indicates that 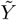 is independent of the features *X* given the response *Y*. Once we swap *X* and *Y*, the conditional independence still holds such that 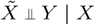. Thus, by swapping *X* and *Y*, model-Y knockoffs satisfy model-X knockoffs in Definition 1.

### 2.2 Algorithm

The proof of Theorem 1 shows that swapping the roles of *X* and *Y* yields valid knockoffs for response variable selection. This fact means that we can switch the roles of *X* and *Y* while fitting a model and perform response selection. In the causal inference context, it is not generally appropriate to swap the features with the responses. However, in the context of response selection, we can swap *X* and *Y* while using model-X knockoffs. Once we swap *X* and *Y*, the original response variables *Y* becomes the features in the swapped model and we follow the three-step procedure of model-X knockoffs described in Section 1.1 to perform the selection. We summarize the key steps in Algorithm 1.

**Algorithm 1.**
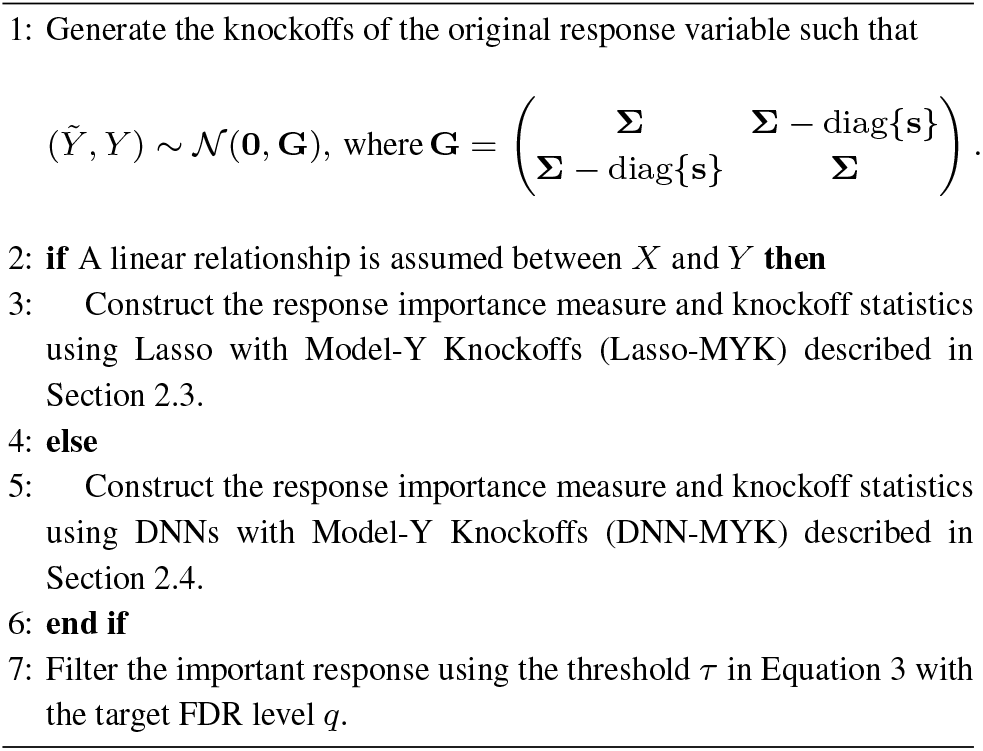
Response Selection using Knockoffs

### 2.3 Generalized Linear Model-Y Knockoffs

To perform response selection under a generalized linear model setting we fit a regularized multinomial logistic regression model using Lasso [Tibshirani, 1996] with the original response *Y* augmented with its knockoffs 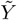 in the role of the covariate variables and the perturbation indicators *X* in the role of the response variable. The optimal Lasso solution is

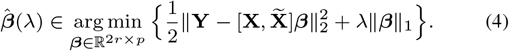

We define 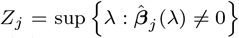 to be the point, *λ*, on the Lasso path at which the response *Y_j_* first enters the model. Then we compute the standard knockoff statistics 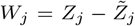, which is likely to be large for positive important responses and negative for null responses. The set of important responses, 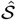, is be selected with controlled FDR using Step 3 as described previously. This Lasso-based procedure can be extended to other generalized linear models.

### 2.4 Nonlinear Model-Y Knockoffs

If a nonlinear relationship between *Y* and *X* is more appropriate for the data set of interest, DNNs can be used to construct the knockoff statistics and perform response selection. But, it is non-trivial to construct a knockoff statistic that satisfies the flip-sign property within a DNN. We introduce a pairwise-competing layer as the first layer of a DNN where the inputs are both the response *Y* and its knockoffs 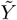. In the pairwise-competing layer, there are *r* nodes and the *j*th node connects only response *Y_j_* and its knockoff 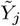 to encourage competition between the two corresponding weights *Ψ_j_* and 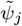. This technique is inspired by Lu et al. [2018] who used it for the purpose of increasing the interpretability and reproducibility of DNNs. We denote the weights from the competing layer to the next layer as *θ*_0_, the weights for the last layer to the output *X* as *θ_L_*, and the weights for each pair of connected intermediate layers as *θ*_1_, *θ*_2_,…, *θ_L_*_−1_. We define Θ_*j*_ = *θ*_0_ ⊙ (*θ*_1_ · *θ*_2_ · · · *θ_L_*), where ⊙ represents entry-wise matrix multiplication. Here, (*θ*_1_ · *θ*_2_ · · · *θ_L_*) represents the matrix multiplication for the weights of all connected layers and can be interpreted as the importance of the *Y_j_* in the multilayer perceptron (MLP). Thus, we define the response importance *Z_j_* as *Ψ_j_* × Θ_*j*_, its knockoff importance 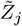 as 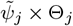 and we obtain the model-Y knockoff statistics 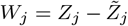, where *j* ∈ [*r*]. As in the case of the Lasso, these definitions are sufficient to employ Theorem 1.

## 3 Synthetic Data Experiments

These synthetic data experiments test whether our theoretical proofs regarding finite-sample fDR control hold empirically. We characterize the response variable selection performance for both linear and nonlinear models by exploring the effects of the key data set properties including: total number of response variables *r*, sample size *n*, correlation *ρ* between features, sparsity level *s* in features.

### Data Generation

We generated synthetic data sets using four mechanisms varying the *model* (linear and nonlinear) and the *features* **X** (continuous or binary) shown in supplementary Table 1. We examined the performance of the model-Y knockoff approach with only binary features in a high-dimensional response selection setting to mimic the setup of our real large-scale genomic data application in Section 4. Table 1 and Table 2 in the Supplementary Information show the overall design of these synthetic data experiments. In the models, **Σ** = CS(*ρ*) is a compound symmetric matrix, with all the diagonal elements being one and all the off diagonal entries being *ρ*. The errors are distributed as 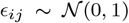. We use (***β***_1_, ***β***_2_,…, ***β***_*r*_) to denote the coefficients vector. The nonlinear model, *g*(*x*) = 2 sin(*x*), is a single-index nonlinear model which has previously been used for benchmark purposes in feature selection procedures using knockoffs [Lu et al., 2018, Zhu and Zhao, 2021]. We denote *m* as the number of important response variables (the number of ***β****_i_* that are nonzero) and define the feature sparsity level *t* as the proportion of the features that do not affect the response in the original (not the swapped) model. For example, if *t* = 0.9, the response variable only depends on 10% of all the features.

**Table 1.** Comparison between marginal testing and knockoffs for LINCS data.

|  | Marginal Only | Knockoffs Only | Both |
| --- | --- | --- | --- |
| Anthracyclines | 0 | 32 | 0 |
| Vorinostat | 0 | 54 | 0 |
| Trichostatin | 0 | 27 | 0 |
| Wortmanin | 0 | 65 | 0 |
| Geldanamycin | 3 | 56 | 12 |
| Sirolimus | 2 | 68 | 2 |

For all synthetic experiments, we set *n* = 400, *r* = 2000, *m* = 10% × *r, ρ* = 0.5, *t* = 0. The nonzero coefficients of ***β***_1_, ***β***_2_,…, ***β***_*r*_ are randomly chosen from {+0.25, −0.25}. To investigate the sensitivity of the power and FDR to key data parameters, we vary one parameter and keep the others at their default level. We summarize the values of each data set parameter in supplementary Table 2.

### Evaluation of FDR and Power

A common approach for variable selection in genomic data applications is to use random forests to identify consistently selected features. Recall, our method for incorporating model-Y knockoffs into a DNN model made use of a pairwise-competing layer. To examine if the pairwise-competing layer is helpful, we adapted the MLP to perform response selection and measured the response importance by the product of the weights for each layer without the pairwise competing layer; we call this method MLP with Model-Y Knockoffs (MLP-MYK). We did not adapt state-of-the-art feature importance learning method for DNNs such as DeepLift [Shrikumar et al., 2017] to response selection since Lu et al. [2018] have shown that DeepLift cannot achieve high power under a high-dimensional setting.

We describe the implementation details of our neural network architecture and tuning parameter choices in the implementation details in the Supplementary Information. In this paper, we try to avoid data set specific tuning as much as possible and choose commonly used parameters for DNNs. Code to reproduce these experiments is available at https://github.com/flahertylab/deepYknock.

### Results and Conclusions

We address four key questions using our simulation experiment data. Can our approach identify the important responses with high power while controlling FDR (1) in either a linear or a nonlinear setting with continuous or binary features? (2) in a high-dimensional setting with *r > n*? (3) when the features are highly correlated? (4) when the responses only rely on part of the features?

We report the power and FDR of our methods Lasso-MYK, DNN-MYK and competing methods RF and MLP-MYK in Figure 1a-1d. We make the following key observations: (1) Both Lasso-MYK and DNN-MYK successfully control FDR across all settings, which validates our theoretical claim. RF fails to control FDR under all settings. MLP-MYK can control FDR but can only achieve dramatically low power, especially in the high dimensional setting with continuous features. (2) Our method Lasso-MYK achieves the highest power under three scenarios: a linear model with continuous features (Figure 1a), a linear model with binary features (Figure 1b), and a nonlinear model with binary features (Figure 1d). (3) Lasso-MYK achieves higher power than the other methods in the challenging settings when the correlation and sparsity level in *X* is high. (4) DNN-MYK achieves higher power than Lasso-MYK under a nonlinear model with continuous features. The performance of our DNN-MYK is much better than MLP-MYK when the feature is continuous under both linear and nonlinear models. This evidence supports the claim that the pairwise-competing layer is useful for identifying important response variables in DNN models.

**Fig. 1:**
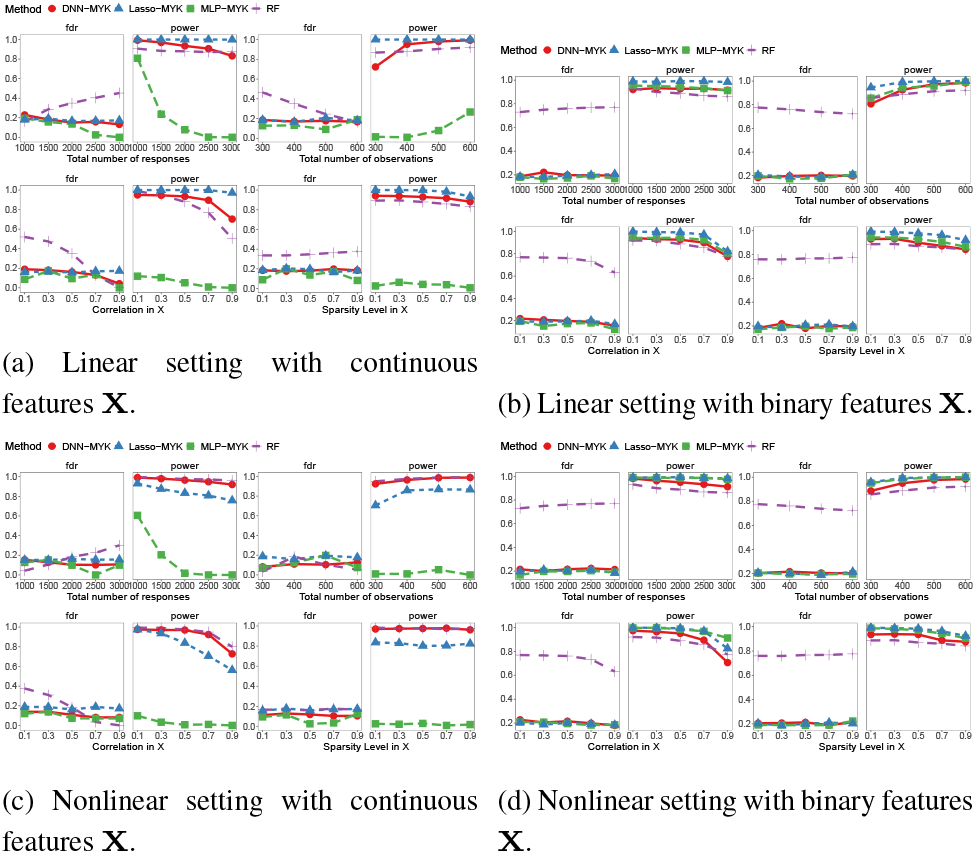
Power and FDR for Lasso-MYK, DNN-MYK, MLP-MYK and Random Forest (RF) across all settings.

## 4 Real Data Analysis

In this study, we examined genomic data from the Library of Integrated Network-Based Cellular Signature (LINCS) Phase I L1000 data set which was collected with the aim to improve understanding of human disease and developing new therapies via cataloging how human cells respond to chemical, genetic and disease perturbations [Keenan et al., 2018]. The data set GSE92742 is fully available online ^1^ and is described in detail elsewhere [Subramanian et al., 2017]. We analyzed a subset of the data dealing with the following small molecule perturbations: anthracycline, vorinostat, trichostatin-a, wortmannin, geldanamycin, sirolimus.

We set the target FDR=0.1 and used Lasso-MYK with binary features in a logistic regression model since it is the most robust method as suggested by our simulation studies. We first present our findings for anthracycline drugs. Then we repeat the same statistical analysis procedure to identify the important genes for vorinostat, trichostatin-a, wortmannin, geldanamycin, and sirolimus, respectively in the Supplementary Information. Finally, we construct a network diagram (Figure 3) to show the individual and shared important transcriptional responses genes when perturbed by the selected six drugs.

### Anthracyclines

For anthracycline drug perturbagens (RUBICIN), the model-Y knockoff procedure selected 40 unique landmark genes across all 100 replications. Figure 2a shows the robust Z-score of the selected genes under DMSO control and anthracycline treatment perturbation. Clearly, the selected important genes have a different signature under the control and treatment groups. In comparison, we also display the average of the knockoffs, 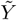, across different replications for the selected genes. The model-Y knockoffs, 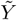, do not demonstrate such strong association with the treatment perturbation. This lack of association is the expected property that we aimed to achieve with 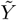 — it mimics the correlation structure in *Y* but is conditionally independent with the perturbation. Figures 2c and 2d show density plot of the robust Z-score and bar plots of the mean and standard error of the coefficients for top 12 selected genes across 100 replications based on their selection frequency.

**Fig. 2:**
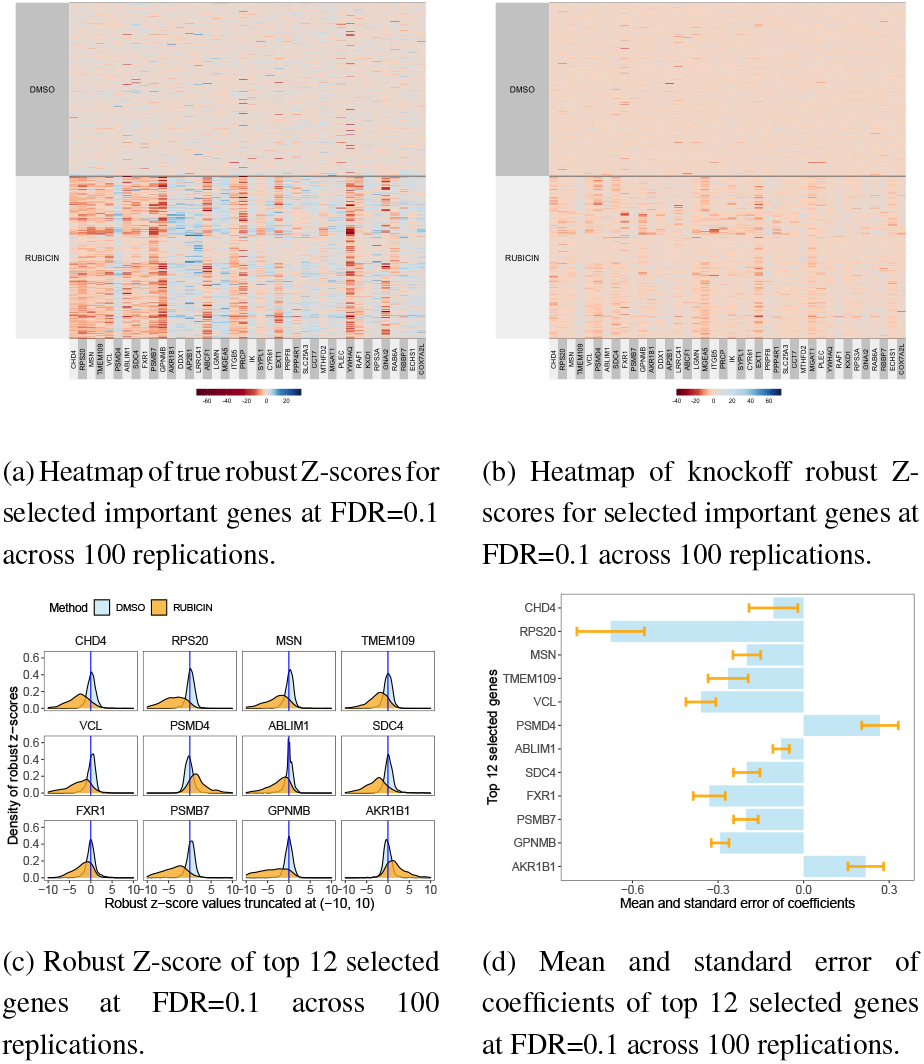
Heatmap of true robust Z-scores for selected genes for anthracycline (a) compared with knockoff responses (b). Density (c) and bar plots (d) with error bars for top selected genes.

Anthracycline compounds are a widely used class of cancer chemotherapy that primarily act by intercalating with DNA and inhibiting both DNA metabolism and RNA synthesis. Lasso-MYK identified Chromodomain Helicase DNA Binding Protein 4 (*CHD4*) which encodes a DNA-binding helicase protein. A recent study reported that “CHD4 depletion dramatically decreases tumor-forming behavior of AML cells and modulates expression of genes associated with tumor colony formation” Sperlazza et al. [2015]. The identification of *CHD4* by our procedure with a negative coefficient suggest that treatment with anthracycline decreases *CHD4* levels and contributes to the anticancer efficacy of the drug by reducing tumor colony formation. Vinculin (*VCL*) is identified as an important response and has a reduced expression after perturbation with anthracycline. Decreased expression of this gene is associated with cardiomyopathy which is a common side effect of anthracycline treatment [Chatterjee et al., 2010]. Finally, FMR1 Autosomal Homolog 1 (*FXR1*) was recently reported in a study involving triple-negative breast cancer patients [Qian et al., 2017]. That study reported that a locus on the q arm of chromosome 3, later localized to *FXR1*, is strongly associated with distant metastasis in triple-negative breast cancer. The observation that *FXR1* is an important gene and has a negative coefficient suggesting the hypothesis that anthracycline may have a therapeutic effect for triple-negative breast cancer patients by decreasing expression of *FXR1* and this reduces the likelihood of distant metastasis.

### Comparison to marginal testing

The standard state-of-the-art methods for identification of important transcriptional responses are based on model-based or model-free marginal testing [Strasser and Weber, 1999, Zeileis et al., 2008]. Therefore, we compare this approach to our conditional testing approach based on knockoffs and to avoid biases due to model selection we select a model-free permutation testing methodology [Strasser and Weber, 1999] and use the local FDR method [Efron, 2004] to control the false discovery rate. We use the `coin` R package to test the independence of the perturbation type and the 978 landmark gene expressions [Zeileis et al., 2008]. Once we obtain the test statistic value for each gene, we used an empirical Bayes technique [Efron, 2004] for large-scale simultaneous hypothesis testing to estimate the local FDR and control the proportion of false positives in the set of genes under the assumption that a majority of the genes modeled are not affected by the perturbations. For each drug, if the estimated FDR for one gene is smaller than the target level of 0.1, the gene will be selected as important. We found that the permutation test method with FDR control at 0.1 is not able to identify any of the genes that were identified by knockoffs for vorinostat, trichostatin-a, wortmannin, and anthracycline, respectively. Table 1 shows an overview of the number of genes identified by marginal testing only, knockoffs only, and both.

For geldanamycin, permutation test with FDR control selects 15 genes including *SEPT2*, *C1D*, *HNRNPK*, *STMN1*, *RPS10*, *RPS21*, *AKR1B1*, *COPB1*, *PPP2CB*, *TPD52L2*, *DHX15*, *COPS6*, *ADAR*, *KDM5A*, *COPS3*. All the selected genes are also identified by our method except *SEPT2*, *RPS21*, and *PPP2CB*. Our method identified 68 out of 978 genes in total under geldanamycin permutations. For sirolimus, permutation test with FDR control only selects *HNRNPK* and *ALDOA*. Both are identified with our method and we identify 70 out of 978 genes in total.

### Response Network

A comparative analysis of the important genes for each small molecule perturbation (Figure 3) reveals the gene expression changes that are private to each perturbation as well as shared genes. Out of five of all six drug perturbations, *CCND2* and *RPS20* are identified as important gene expressions that are affected by the drugs. *CCND2* has been identified as a common therapeutic target in lung and breast cancer [Hung et al., 2018]. Our identification of the transcriptional response to perturbations by all compounds is interesting in that it may indicate a wider molecular role for this gene. Diseases associated with *RPS20* include familial colorectal cancer type X and diamond-blackfan anemia. *RPS20* shown in Figure 3 has a negative coefficient with all six drugs which indicates a decrease in *RPS20* levels which in turn suggests the potential to contribute to treatment of anti-familial colorectal cancer and diamond-blackfan anemia.

**Fig. 3:**
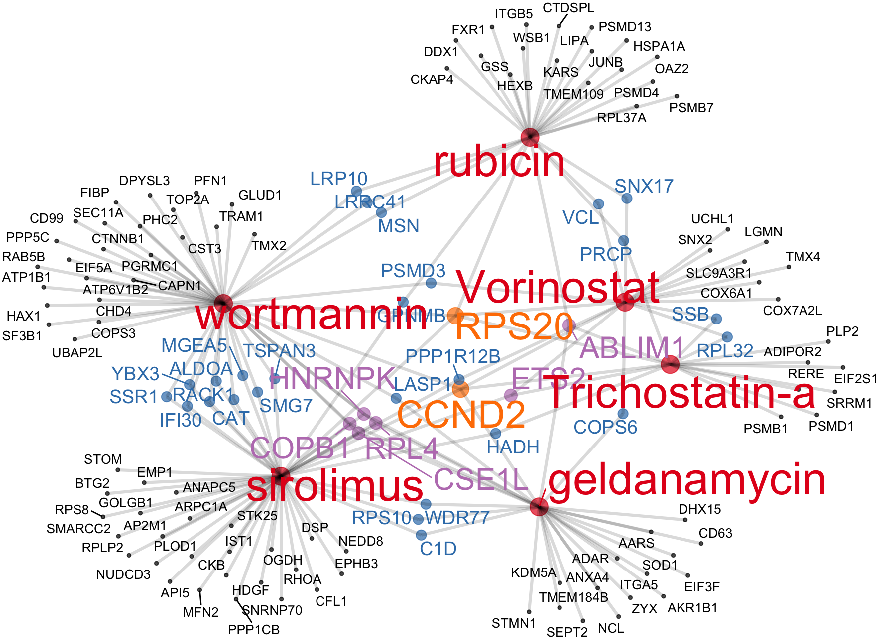
Network plot of identified genes for vorinostat, trichostatin-a, wortmannin, geldanamycin, sirolimus, and rubicin. Genes in orange, purple, blue, and black represent the common genes identified with exposure to five, three, two and one drug, respectively.

Heterogeneous Nuclear Ribonucleoprotein K (*HNRNPK*) was identified as important and has a negative coefficient under wortmannin, geldanamycin, sirolimus perturbations (supplementary Figure 4, 5, 6). Poenisch et al. [2015] reported that an RNA interference of *HNRNPK* results in decreased Hepatitis C virus (HCV) particle production without affecting viral RNA replication. Therefore, our findings suggest that wortmannin, geldanamycin, sirolimus may decreased *HNRNPK* expression and thus decrease HCV particle production. In line with this hypothesis, wortmannin has been suggested as pretreatment drugs for acute liver damage [Li et al., 2014].

Human chromosomal segregation 1-like (*CSE1L*) expression is associated with tumor progression in various human cancers. Li et al. [2020] showed that *CSE1L* was highly expressed in gastric cancer cell lines and *CSE1L* silencing promoted apoptosis and inhibited cell proliferation and invasion. We have found that the the coefficient of *CSE1L* is negative under vorinostat, wortmannin and geldanamycin perturbations (Supplementary Figures 2, 4, 5), which indicates that vorinostat, wortmannin and geldanamycin are able to decrease expressions of *CSE1L* which may indicate its benefits for gastric cancer treatment.

## 5 Discussion

In this paper we have presented a novel method for controlled response variable selection with theoretical finite sample guarantees on the FDR. We show how the framework can be used in both linear and nonlinear (DNN) models. Synthetic and real data analysis of experimental genomics data sets empirically support the theoretical behavior of the approach. Our analysis of the NIH LINCS data identified important genes whose expression is modulated in response to perturbation with anthracycline, vorinostat, trichostatin-a, geldanamycin, and sirolimus, small molecule drugs used in cancer chemotherapy using NIH LINCS data set. Furthermore, we compared the set of important genes that respond to vorinostat, geldanamycin, sirolimus, trichostatin-a, wortmannin, and anthracycline to elucidate the response gene network and identify co-responsive pathways. A *potential limitation* is that we only consider a Gaussian distribution of the response variable such that the *Y* knockoffs can be generated analytically according Equation (1). Even so, we have found that the Gaussian assumption practically useful in a real data analysis scenario and the knockoff framework has been shown to be robust to this assumption on *F_X_* [Barber et al., 2018]. There are several areas that are interesting for future work. In this work, we have assumed the marginal distribution *F_Y_* is known, and this assumption has proven to be reasonable for the data set we analyzed. But in other applications, it may be necessary to learn about the marginal distribution and it would be of interest to offer theoretical bounds on the performance when approximations are employed. In addition, it is also of great interest to design more flexible generation methods to relax the Gaussian assumption for model-Y knockoffs.

## Funding

This work was supported by NSF Award 1934846 and NIH R01-GM135931.

## Data Availability

The data set GSE92742 is fully available online ^2^ and is described in detail elsewhere Subramanian et al. [2017].

## Footnotes

1 http://lincsportal.ccs.miami.edu/dcic-portal/

2 http://lincsportal.ccs.miami.edu/dcic-portal/

## Notes

https://github.com/flahertylab/deepYknockoff

https://www.ncbi.nlm.nih.gov/geo/query/acc.cgi?acc=GSE92742

### Competing Interest Statement

The authors have declared no competing interest.

