## Supplemental Information for "Identification of Significant Gene Expression Changes in Multiple Perturbation Experiments using Knockoffs"

### A Synthetic Data Experiment Setup

We generated synthetic data sets using four mechanisms varying the *model* (linear and nonlinear) and the *features* $\mathbf{X}$ (continuous or binary) shown in Table 1. Table 1 and Table 2 show the overall design of these synthetic data experiments. In the models, $\mathbf{\Sigma} = \mathrm{CS}(\rho)$ is a compound symmetric matrix, with all the diagonal elements being one and all the off diagonal entries being $\rho$. The errors are distributed as $\epsilon_{ij} \sim \mathcal{N}(0, 1)$. We use $(\boldsymbol{\beta}_1, \boldsymbol{\beta}_2, \ldots, \boldsymbol{\beta}_r)$ to denote the coefficients vector. The nonlinear model, $g(x) = 2\sin(x)$, is a single-index nonlinear model which has previously been used for benchmark purposes in feature selection procedures using knockoffs [Lu et al., 2018, Zhu and Zhao, 2021].

| Models | Continuous Features | Binary Features |
|---|---|---|
| Linear | Each row of $\mathbf{X} \sim \mathcal{N}(\mathbf{0}, \mathbf{\Sigma})$ | Each row of $\mathbf{X} \sim \mathcal{N}(\mathbf{0}, \mathbf{\Sigma})$ |
| | $\mathbf{Y}_{n\times r} = \mathbf{X}_{n\times p}\boldsymbol{\beta} + \boldsymbol{\epsilon}_{n\times r}$ | $\mathbf{Y}_{n\times r} = \mathbf{X}_{n\times p}\boldsymbol{\beta} + \boldsymbol{\epsilon}_{n\times r}$ |
| | | $\mathbf{X}^* = \mathbf{1}(\mathbf{X} > 0)$ |
| | Observe $(\mathbf{Y}_{n\times r},\ \mathbf{X}_{n\times p})$ | Observe $(\mathbf{Y}_{n\times r},\ \mathbf{X}^*_{n\times p})$ |
| Nonlinear | Each row of $\mathbf{X} \sim \mathcal{N}(\mathbf{0}, \mathbf{\Sigma})$ | Each row of $\mathbf{X} \sim \mathcal{N}(\mathbf{0}, \mathbf{\Sigma})$ |
| | $\mathbf{Y}_{n\times r} = 2\sin(\mathbf{X}_{n\times p}\boldsymbol{\beta}) + \boldsymbol{\epsilon}_{n\times r}$ | $\mathbf{Y}_{n\times r} = 2\sin(\mathbf{X}_{n\times p}\boldsymbol{\beta}) + \boldsymbol{\epsilon}_{n\times r}$ |
| | | $\mathbf{X}^* = \mathbf{1}(\mathbf{X} > 0)$ |
| | Observe $(\mathbf{Y}_{n\times r},\ \mathbf{X}_{n\times p})$ | Observe $(\mathbf{Y}_{n\times r},\ \mathbf{X}^*_{n\times p})$ |

Table 1: Synthetic data generation mechanisms from each combination of model and feature type, where $\boldsymbol{\beta} = (\boldsymbol{\beta}_1, \boldsymbol{\beta}_2, \ldots, \boldsymbol{\beta}_r)$.

| **Parameter** | **Levels** |
|---|---|
| Number of responses ($r$) | $1000, 1500, 2000, 2500, 3000$ |
| Sample size ($n$) | $300, 400, 500, 600$ |
| Correlation ($\rho$) in $\mathbf{X}$ | $0.1, 0.3, 0.5, 0.7, 0.9$ |
| Sparsity level ($t$) in $\boldsymbol{\beta}$ | $0.1, 0.3, 0.5, 0.7, 0.9$ |

Table 2: Values of the key parameter values in each experiment setting.

### B Implementation Details

The number of nodes in each layer of our neural network is $2r - r - 64 - 32 - p$ from the input to the output, where $r$ denotes the total number of response variables and $p$ denotes the number of features. The first layer is a pairwise-competing layer and the rest layers are fully connected. In the first pairwise-competing layer, there are $r$ nodes and the $j$th node connects only response $Y_j$ and its knockoff $\tilde{Y}_j$ to encourage competition between the two corresponding weights

$\psi_j$ and $\tilde{\psi}_j$. L1-regularization and L2-regularization are used in all layers and the tuning parameters are chosen as 0.001 for both. ReLU [Nair and Hinton, 2010] activation are only used for the hidden layers. Adam optimizer [Kingma and Ba, 2014] is used as the optimization engine to update the neural network. In this paper, we try to avoid data set specific tuning as much as possible. We choose commonly used parameters for deep neural networks (DNNs). The batch size is set as 100 and the learning rate is set as 0.001 with a common decay rate of 1e-5 for every epoch in a batch setting. The number of epochs is 100. All numerical experiments are run on a MacBook Pro with a 2.4 GHz Quad-Core Intel Core i5 processor and 16GB memory. The algorithm is implemented in Python 3.8.5.

### C Real Data Preprocessing

There are five data processing levels in the NIH LINCS L1000 data set; we focus on the level 4 data which provides a robust Z-scores for each gene normalized with respect to the population of vehicle controls and the entire plate population. The robust Z-score is the median absolute deviation (MAD) of a data set and is more robust to single outliers than the mean. The score is computed as: robust Z-score $= (x - \mu_{1/2})/\text{MAD}$, where MAD is defined as the median of the absolute deviation from the median of the data points $X_1$ to $X_n$: $\text{MAD} = \text{median}_i(\mid X_i - \text{median}(X_1, X_2, \ldots, X_n) \mid)$. We took a subset of the data to focus on small molecule compounds including: vorinostat, trichostatin-a, wortmannin, geldanamycin, sirolimus, all anthracycline drug perturbagens (RUBICIN) including "daunorubicin", "epirubicin", "idarubicin", "doxorubicin", and "pirarubicin" giving a total treatment sample size of $n = 12,247$. We obtain the control group by randomly sampling $n = 12,247$ samples with Dimethylsulfoxide (DMSO) perturbations since DMSO is the control for compound treatments considered in L1000.

We analyzed a subset of the data dealing with the following small molecule perturbations:

**vorinostat:** a class of drugs in chemotherapy that can treat cutaneous T-cell lymphoma.

**trichostatin-a:** an organic compound that serves as an antifungal antibiotic and has some potential as an anti-cancer drug.

**wortmannin:** inhibits growth of cancer [Liu et al., 2005], inhibits proliferation, induces apoptosis [Kim et al., 2012] and promotes cell death [Mizuno et al., 2003, Jiang et al., 2011].

**geldanamycin:** an antitumor antibiotic that inhibits the function of Hsp90 by binding to the unusual ADP/ATP-binding pocket of the protein, where HSP90 client proteins play important roles in the regulation of the cell cycle, cell growth, cell survival, apoptosis, angiogenesis and oncogenesis.

**sirolimus:** an immunosuppressive drug that can prevent organ rejection after a kidney transplant.

**anthracycline:** a class of drugs in cancer chemotherapy.

For cost efficiency, a custom L1000 instrument was developed for the project to directly measure $r = 978$ "landmark" genes and to enable measurement of millions of perturbation samples. Landmark genes were selected to make imputation of other genes possible. We found that the empirical pairwise correlation between landmark genes is below 0.5 for almost all pairs, indicating little multicollinearity in the marginal distribution of the responses. We show the histogram of pairwise correlation between all pairs of 978 landmark genes in Figure 1.

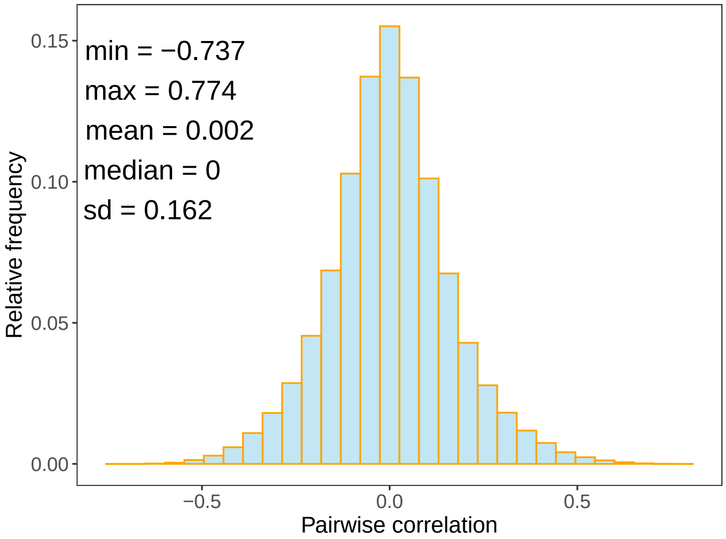


Figure 1: Histogram of pairwise correlation between all pairs of 978 landmark genes.

After preprocessing, a response matrix of dimension $n \times r$ is obtained, where $n = 24,494$ and $r = 978$. The entries of the response matrix contain the robust Z-scores for a gene (column) after perturbation (row). We divided this response matrix into six different subsets where each subset only gets exposure to one of the six drugs. The feature or covariate matrix is a single binary vector indicating whether the perturbation is a control or a treatment with vorinostat, trichostatin-a, wortmannin, geldanamycin, sirolimus, or anthracycline drugs. Like other knockoff generation methods, our method is not deterministic, and the selected response variables depend on the realization of $\tilde{Y}$ in each replication Sesia et al. [2019]. Thus, we report a ranking of these results over 100 independent replications of the model-Y knockoffs procedure.

#### D Real Data Analysis Results

**Vorinostat.** The negative coefficient of *COXA1* indicates that vorinostat has treatment effect on Hurthle cell neoplasms (Figure 2). It has been reported that Cytochrome C Oxidase Subunit 6A1 (*COXA1*) is involved in mitochondrial metabolism and is significantly overexpressed in tumors in Hurthle cell neoplasms versus normal tissue [Canberk et al., 2021]. D'Arcy and Linder [2014] reported that Ubiquitin C-Terminal Hydrolase L1 (*UCHL1*) showed decreased activity in the tumor tissues, potentially indicating a tumor-suppressive role for this protein. It has also been reported that loss of *UCHL1* expression is a common occurrence with tumor tissues and silencing via methylation of the *UCHL1* promoter has also been observed in biopsies of pancreatic cancer [Pérez-Mancera et al., 2012] and prostate cancer [Sato et al., 2003, Ummanni et al., 2011]. The positive coefficient of *UCHL1* (Figure 2) in our analysis is concordant with these previous observations.

**Trichostatin-a.** Our regression analysis produced a positive coefficient (Figure 3) for Cyclin D2 (*CCND2*) which indicates that *CCND2* expression increases with treatment with Trichostatin-a. Takai et al. [2004] have shown that Histone deacetylase inhibitors (HDACIs) [suberoyl anilide bishydroxamine, valproic acid (VPA), trichostatin A, and sodium butyrate] are able to inhibit endometrial cancer cell proliferation and stimulate apoptosis. In the altered gene expressions related to the malignant phenotype, there is a decrease in Cyclin D1 (*CCND1*) and *CCND2*. The positive coefficient of *CCND2* indicates a potential beneficial treatment effect of trichostatin A.

**Wortmannin.** It has been reported that a single nucleotide polymorphism in Meningioma expressed antigen 5 (*MGEA5*), encoding O-Glcnacase (OGA), is associated with type 2 diabetes in a Mexican American population [Lehman et al., 2005]. Wortmannin also has been found to have an effect on reducing insulin signaling and death in seizure-prone mice [MacKay et al., 2012]. In our study, we find that the expression of *MGEA5* exposed to wortmannin perturbations is increased (Figure 4). This leads to the hypothesis that the genetic polymorphism leads to decreased protein activity and treatment by Wortmannin reverses the effects of this decrease by increasing expression levels of the protein.

**Geldanamycin.** Hung et al. [2018] reported that *CCND2* is a common biomarker in lung and breast cancer. They suggested that increased *CCND2* expression can help inhibit cancer cell growth and migration ability in Taiwanese breast cancer patients. In our study, the positive coefficient (Figure 5) estimate of *CCND2* indicates that the exposure of geldanamycin can help increase expression of *CCND2* and geldanamycin can be a potential drug for breast cancer and lung cancer and is consistent with preliminary reports of its potential [Ochel et al., 2001].

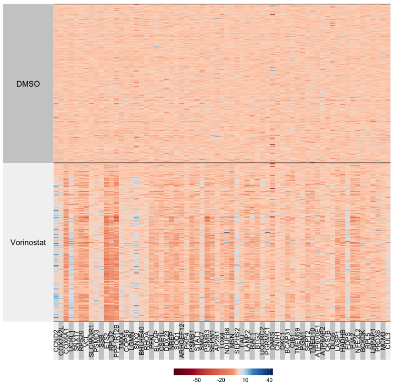


(a) Heatmap of true robust Z-scores for selected important genes at FDR=0.1 across 100 replications.

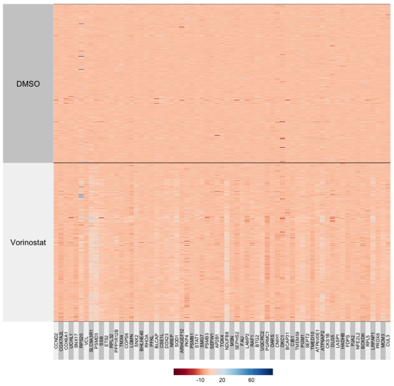


(b) Heatmap of knockoff robust Z-scores for selected important genes at FDR=0.1 across 100 replications.

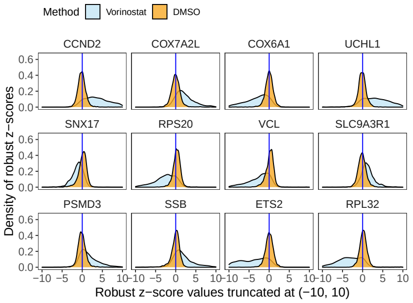


(c) Robust Z-score of top 12 selected genes at FDR=0.1 across 100 replications.

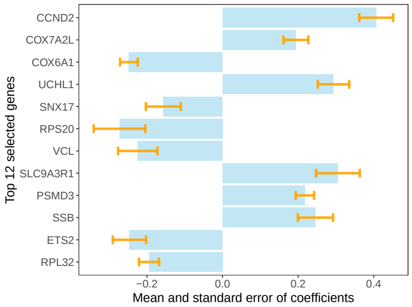


(d) Mean and standard error of coefficients of top 12 selected genes at FDR=0.1 across 100 replications.

Figure 2: Heatmap of true robust Z-scores for selected genes for vorinostat (a) compared with knockoff responses (b). Density (c) and bar plots (d) with error bars for top selected genes.

**Sirolimus.** Glycoprotein Nmb (*GPNMB*) is a melanoma-associated glycoprotein that is targeted by the CR011-vcMMAE antibody-drug conjugate (ADC) and it has been shown that CR011-vcMMAE induces the apoptosis of GPNMB-expressing tumor cells in vitro and tumor regression in xenograft models [Qian et al., 2008]. That report also suggested the possibility of increasing the anticancer activity of CR011-vcMMAE through pharmacological enhancement of *GPNMB* expression for potential therapeutic benefit. The positive coefficient ( Figure 6) of *GPNMB* indicates that sirolimus indeed may have potential therapeutic benefit in combination with other anticancer drugs.

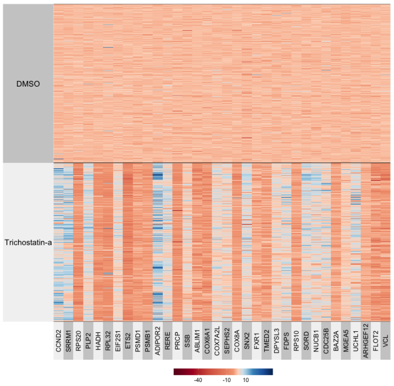


(a) Heatmap of true robust Z-scores for selected important genes at FDR=0.1 across 100 replications.

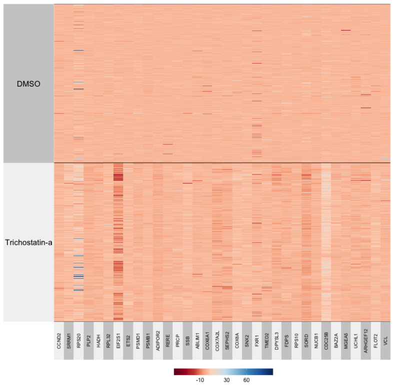


(b) Heatmap of knockoff robust Z-scores for selected important genes at FDR=0.1 across 100 replications.

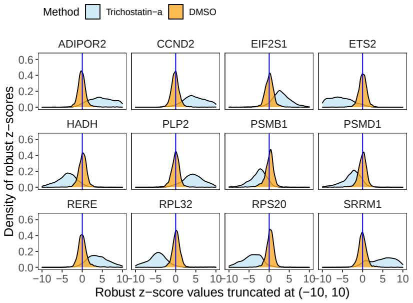


(c) Robust Z-score of top 12 selected genes at FDR=0.1 across 100 replications.

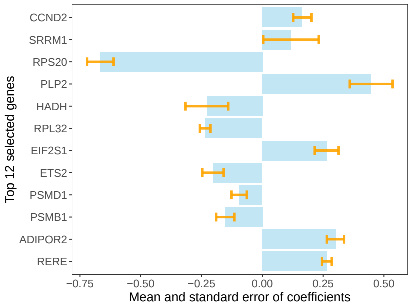


(d) Mean and standard error of coefficients of top 12 selected genes at FDR=0.1 across 100 replications.

Figure 3: Heatmap of true robust Z-scores for selected genes for trichostatin-a (a) compared with knockoff responses (b). Density (c) and bar plots (d) with error bars for top selected genes.

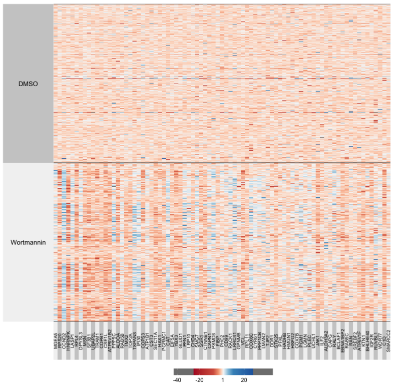


(a) Heatmap of true robust Z-score for selected important genes at FDR=0.1 across 100 replications.

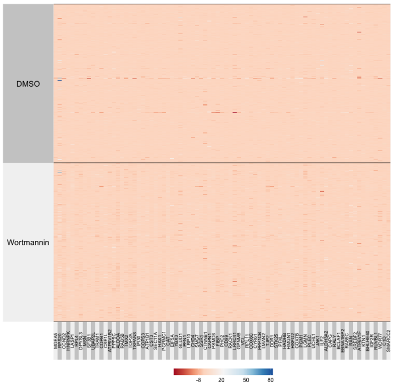


(b) Heatmap of knockoff robust Z-score for selected important genes at FDR=0.1 across 100 replications.

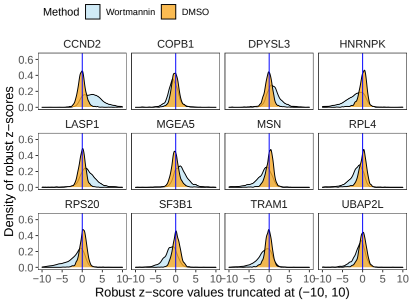


(c) Robust Z-score of top 12 selected genes at FDR=0.1 across 100 replications.

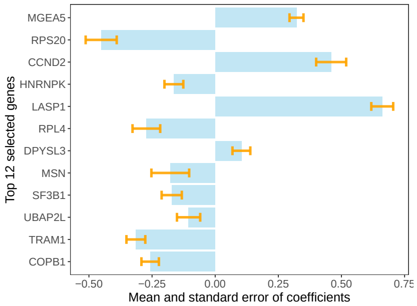


(d) Mean and standard error of coefficients of top 12 selected genes at FDR=0.1 across 100 replications.

Figure 4: Heatmap of true robust Z-scores for selected genes for wortmannin (a) compared with knockoff responses (b). Density (c) and bar plots (d) with error bars for top selected genes.

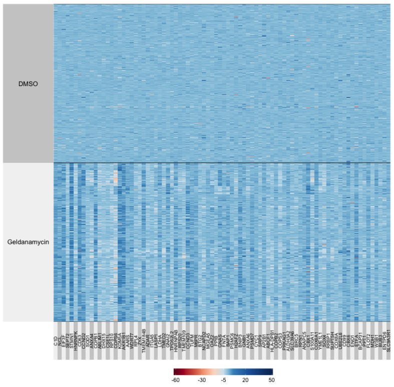


(a) Heatmap of true robust Z-score for selected important genes at FDR=0.1 across 100 replications.

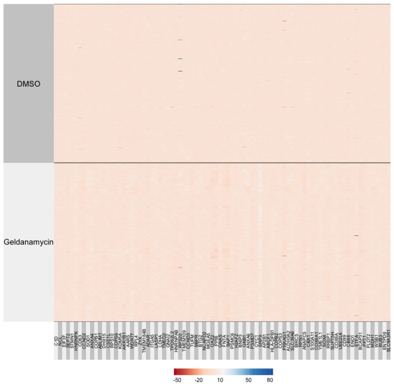


(b) Heatmap of knockoff robust Z-score for selected important genes at FDR=0.1 across 100 replications.

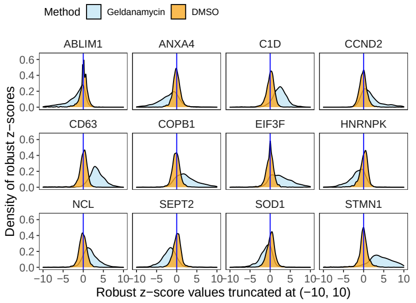


(c) Robust Z-score of top 12 selected genes at FDR=0.1 across 100 replications.

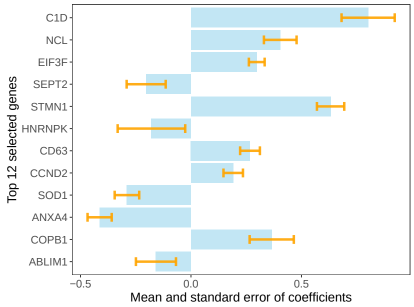


(d) Mean and standard error of coefficients of top 12 selected genes at FDR=0.1 across 100 replications.

Figure 5: Heatmap of true robust Z-scores for selected genes for geldanamycin (a) compared with knockoff responses (b). Density (c) and bar plots (d) with error bars for top selected genes.

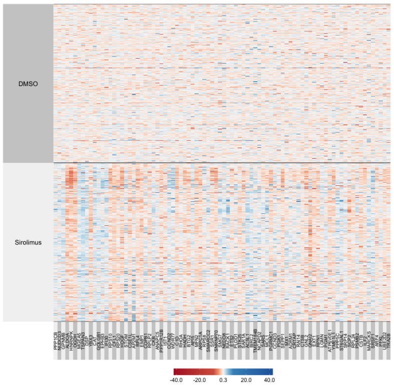


(a) Heatmap of true robust Z-score for selected important genes at FDR=0.1 across 100 replications.

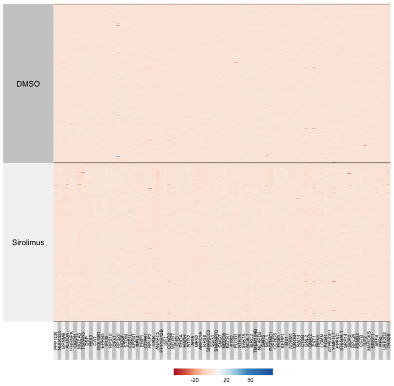


(b) Heatmap of knockoff robust Z-score for selected important genes at FDR=0.1 across 100 replications.

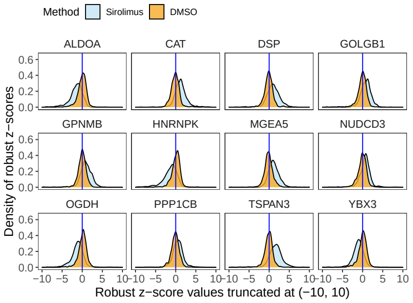


(c) Robust Z-score of top 12 selected genes at FDR=0.1 across 100 replications.

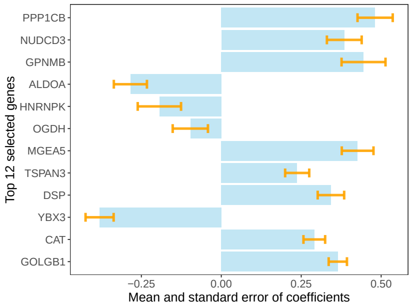


(d) Mean and standard error of coefficients of top 12 selected genes at FDR=0.1 across 100 replications.

Figure 6: Heatmap of true robust Z-scores for selected genes for sirolimus (a) compared with knockoff responses (b). Density (c) and bar plots (d) with error bars for top selected genes.
